## Supplementary figures and images for "VGLL2 and TEAD1 fusion proteins drive YAP/TAZ-independent tumorigenesis by engaging p300"

### Figure 1-figure supplement 1

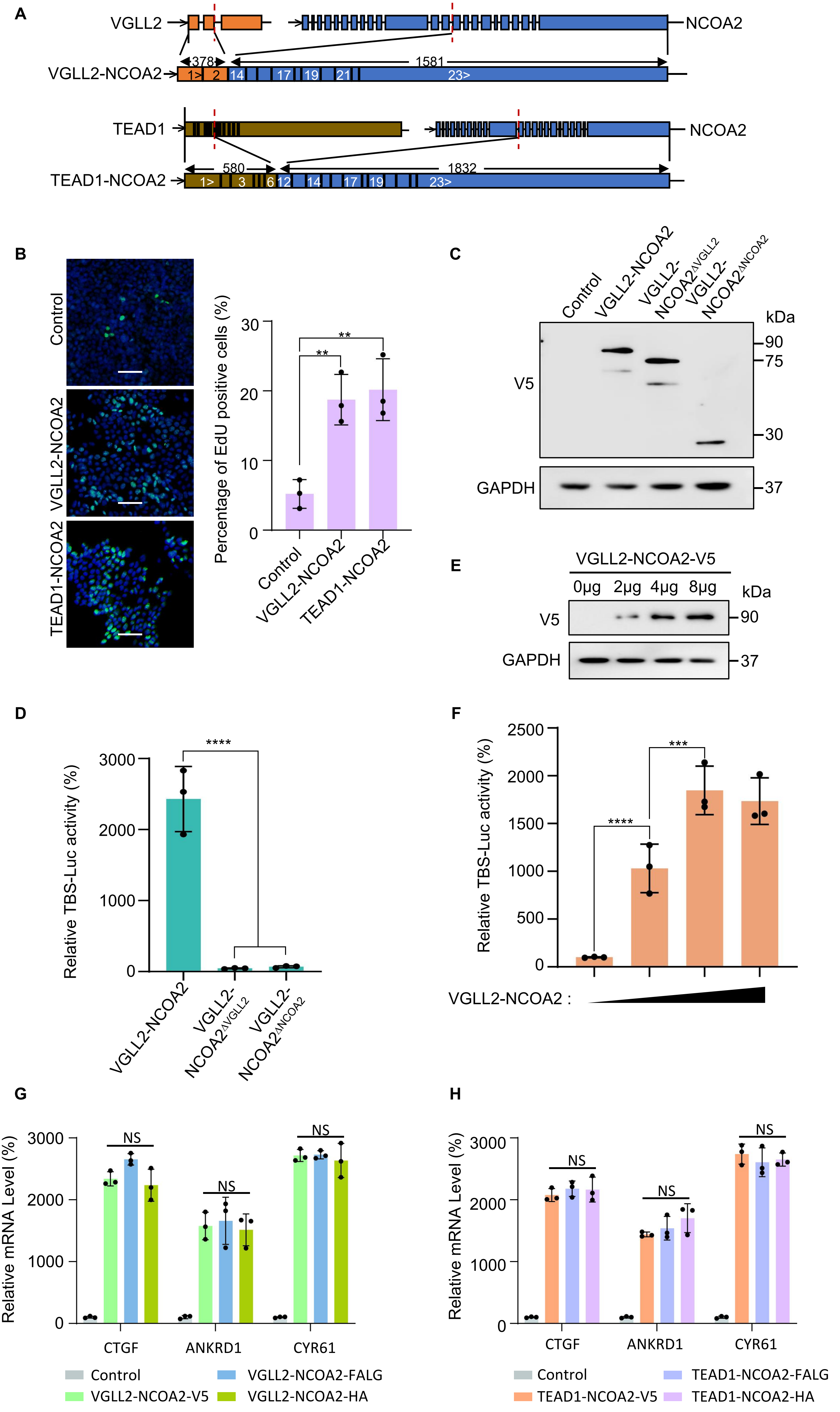

### Figure 3-figure supplement 1

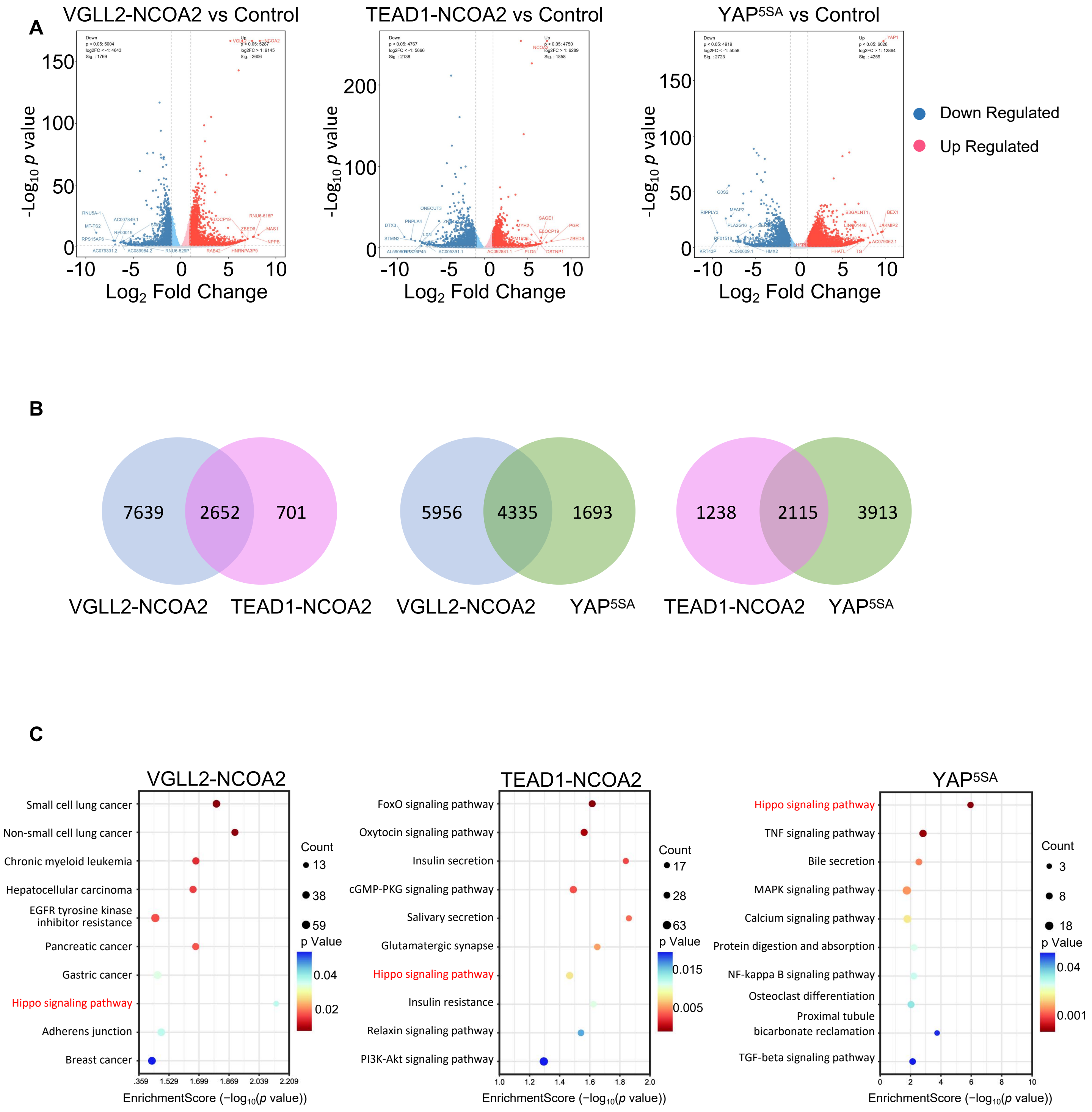

### Figure 3-figure supplement 2

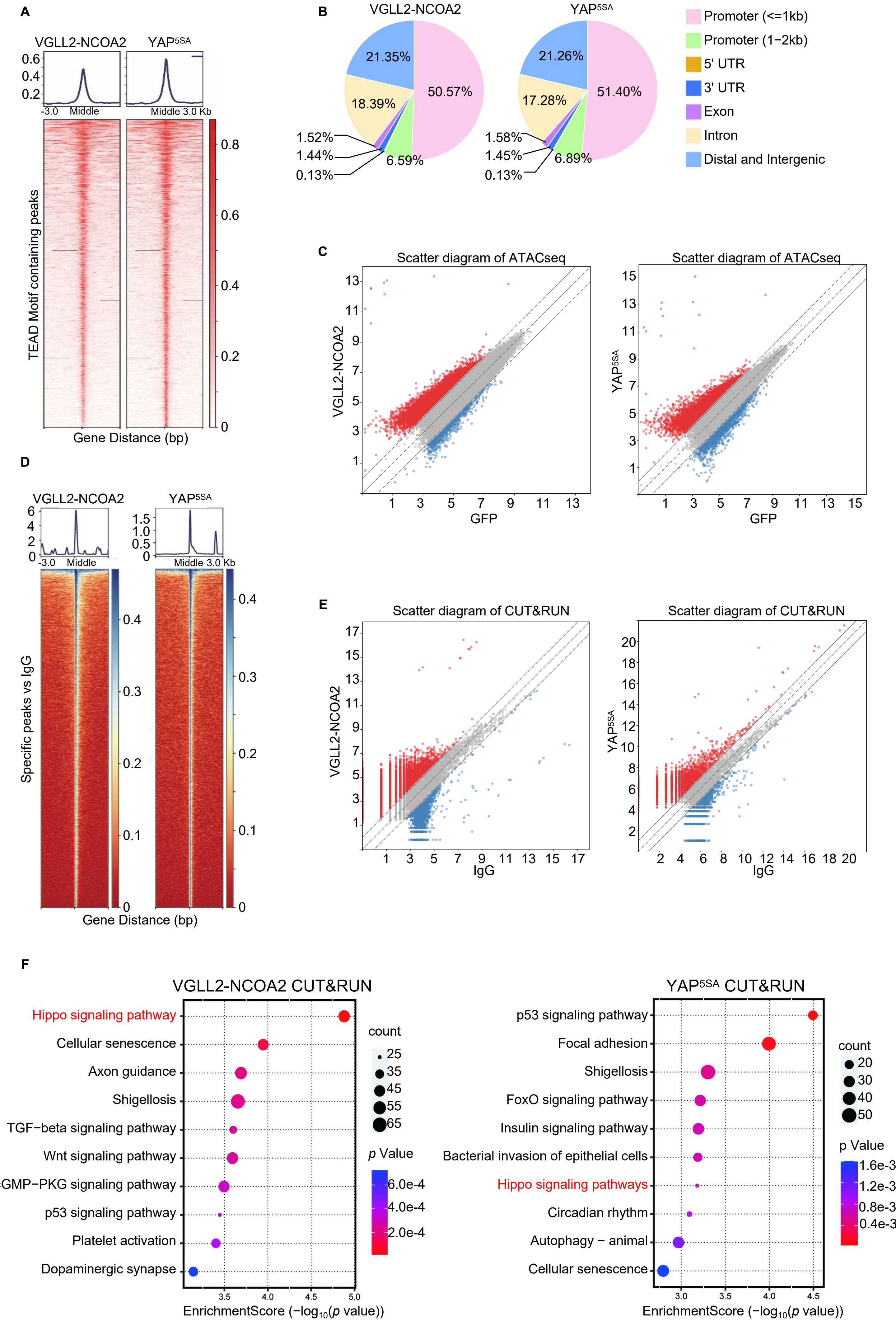
